# Genetic tools for the stable overexpression of circular RNAs

**DOI:** 10.1101/2021.05.27.446018

**Authors:** Nicol Mecozzi, Arianna Nenci, Olga Vera, Aimee Falzone, Gina M. DeNicola, Florian A. Karreth

**Author notes:** Corresponding Author: Florian A. Karreth.

## Abstract

Circular RNAs (circRNAs) are a class of non-coding RNAs that feature a covalently closed ring structure formed through backsplicing. circRNAs are broadly expressed and contribute to biological processes through a variety of functions. Standard gain-of-function and loss-of-function approaches to study gene functions have significant limitations when studying circRNAs. Overexpression studies in particular suffer from the lack of efficient genetic tools. While mammalian expression plasmids enable transient overexpression of circRNAs in cultured cells, most cell biological studies require long-term ectopic expression. Here we report the development and characterization of genetic tools enabling stable circRNA overexpression *in vitro* and *in vivo*. We demonstrated that circRNA expression constructs can be delivered to cultured cells via transposons, whereas lentiviral vectors have limited utility for the delivery of circRNA constructs. We further showed that circRNA transposons can be supplied to mouse livers via hydrodynamic tail vein injection, resulting in ectopic circRNA expression in a hepatocellular carcinoma mouse model. Furthermore, we generated genetically engineered mice harboring circRNA expression constructs. We demonstrate that this approach enables constitutive, global circRNA overexpression as well as inducible circRNA expression directed specifically to melanocytes in a melanoma mouse model. Overall, these tools expand the genetic toolkit available for the functional characterization of circRNAs of interest.

## Introduction

The last decade of RNA research revealed a myriad of non-coding RNAs that support fundamental biological processes and whose deregulation may contribute to the development of diseases such as cancer. Circular RNAs (circRNAs) are among the most recent additions to the non-coding RNA compendium and represent a peculiar class of RNA molecules. Generated from pre-mRNA via a process termed backsplicing where the 3’ end of a downstream exon is spliced to the 5’ end of an upstream exon, circRNAs form covalently closed ring molecules lacking 5’ CAP and 3’ polyA tail features. The circular structure protects also from exonuclease cleavage, rendering circRNAs extraordinarily stable. Advances in RNA sequencing led to the identification of thousands of circRNAs in humans and other species. Interestingly, circRNA expression is cell type- or differentiation state-dependent and often uncoupled from the expression of their cognate linear RNAs from which they derive^1–10^. This suggests that circRNA formation is a regulated process and that circRNAs must serve important functions in the cell. Indeed, circRNAs have been shown to act as natural microRNA sponges^3,11^, associate with RNA-binding proteins^12–16^ and chromatin^17^, and regulate the transcription of their parental genes^18–20^. Some circRNAs even contain internal ribosome entry site-like structures and encode proteins^21–26^. Thus, circRNA functions are diverse and unpredictable based on their sequence alone, requiring functional characterization to reveal the underlying mode of action.

Functional analyses of circRNAs have been hindered by technical challenges. Knock-down experiments using RNAi require that the siRNA or shRNA is targeted to the backsplice junction to avoid simultaneous silencing of the cognate linear mRNA. This limits the possibilities for designing potent siRNA/shRNA and may lead to off-target effects on the cognate linear mRNA or other transcripts. Recently, a CRISPR approach using an engineered type VI-D Cas13d enzyme, CasRx^27^, demonstrated superior circRNA silencing in terms of efficiency and specificity^28^. The overexpression of a circRNA of interest may be even more difficult than silencing. Circularization requires the presence of flanking introns containing repeat elements, which upon their alignment bring the appropriate splice sites into close proximity to aid in the backsplicing reaction. Mammalian expression plasmids have been generated where circRNA exons are flanked by introns derived from genes that efficiently produce circRNAs, for instance the *D. melanogaster* Laccase2 introns or the human ZKSCAN1 introns^29,30^. With these expression plasmids efficient ectopic circRNA expression can be achieved in mammalian or *D. melanogaster* cells^29^. However, such plasmids do not integrate into the genome and ectopic circRNA expression is therefore only temporary. This significantly limits long-term phenotypic analyses of cell biological effects. In addition, whether similar approaches are amenable to circRNA overexpression in genetically engineered mice for functional circRNA studies *in vivo* is not known.

Here, we tested stable, long-term circRNA overexpression approaches. While lentiviral circRNA delivery performed poorly, we found that using a transposon for delivery and genomic integration of circRNA expression constructs enables long-term circRNA expression in cells and a hepatocellular carcinoma mouse model. Moreover, we generated genetically engineered mice that overexpress a circRNA either constitutively and ubiquitously or inducibly in a cell-type specific manner in a melanoma model. These tools and approaches will enable more in-depth analyses of circRNA functions *in vitro* and *in vivo*.

## Results

### Limited utility of lentiviruses for circRNA overexpression

To identify efficient approaches that enable long-term, stable expression of ectopic circRNAs, we first tested the utility of lentiviruses. To this end, we cloned a split-GFP (spGFP) reporter into the pLenti-Blasticidin lentiviral vector under the control of the CMV promoter (spGFP-pLenti). The spGFP reporter^29^ contains a split enhanced GFP cDNA and an internal ribosome entry site (IRES) which allows for the production of GFP protein from the circRNA. Circularization is mediated by the introns flanking the human ZKSCAN1 circRNA (Fig. 1A). We transfected the spGFP-pLenti vector along with VSVG and Δ8.2 packaging and envelope plasmids into HEK293 cells for lentivirus production. We also transfected a pLenti-Blasticidin version carrying linear GFP (linGFP-pLenti, Fig. 1A) into HEK293 cells for side-by-side comparison of lentiviral vectors carrying constructs that are spliced (spGFP) or not spliced (linGFP). linGFP-pLenti-transfected HEK293 cells expressed GFP (Fig. 1B), which is expected due to translation of the viral transcript. Notably, HEK293 cells transfected with spGFP-pLenti also were GFP-positive, albeit at lower levels (Fig. 1B). In addition, circularized spGFP RNA was detected in spGFP-pLenti-transfected HEK293 cells by qRT-PCR using divergent primers that specifically detect backspliced RNA (Fig. 1C). Circular spGFP was not detected in linGFP-pLenti-transfected HEK293 cells, while using convergent primers that do not distinguish between circular and linear transcripts detected GFP in spGFP-pLenti- and linGFP-pLenti-transfected cells (Fig. 1C). We also detected GFP protein in spGFP-pLenti-transfected HEK293 cells by Western blot (Fig. 1D). This indicates that the spGFP-pLenti viral transcript is backspliced prior to packaging into viral capsids, which may interfere with efficient virus production. To test this, we first estimated the viral titer by measuring the amount of p24 protein in the viral supernatant using the Lenti-X GoStix kit. The viral titer produced by spGFP-pLenti was even greater than that produced by linGFP-pLenti (Fig. 1E), indicating that backsplicing does not impair the generation of viral particles. We then infected two human melanoma cell lines, WM164 and 501Mel, with the spGFP and linGFP lentiviral particles and selected the cells in 10µg/mL Blasticidin for 10 days. Notably, despite the greater amount of p24 protein in the spGFP-pLenti viral supernatant, melanoma cells infected with this virus preparation died significantly more during Blasticidin selection than cells infected with linGFP-pLenti (Fig. 1F). WM164 and 501Mel cells infected with spGFP-pLenti that survived Blasticidin selection were negative for GFP by fluorescence imaging (Fig. 1G). Moreover, we could not detect circularized spGFP by qRT-PCR using divergent primers and only very minor amounts of GFP using convergent primers that detect circularized and non-circularized GFP (Fig. 1H). Overall, these findings suggest that lentiviruses may be suboptimal for the generation of stable cell lines overexpressing circRNAs.

**Figure 1:**
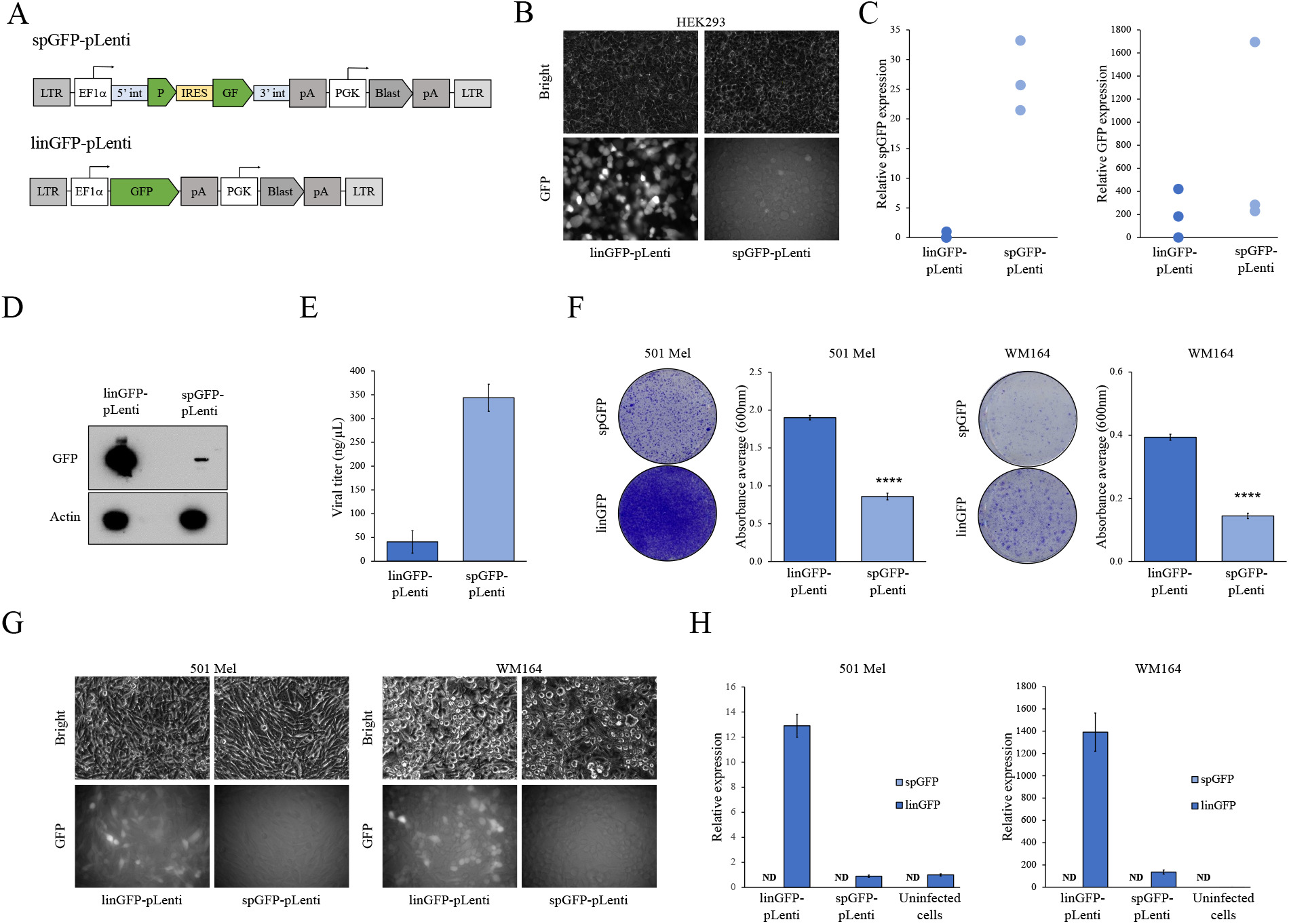
Lentiviral delivery of a circRNA expression construct. (A) Schematic outline of the circular spGFP (spGFP-pLenti) and linear GFP (linGFP-pLenti) expression constructs. LTR, long-terminal repeat; 5’int, human ZKSCAN1 5’ intron; 3’int, human ZKSCAN1 3’ intron; IRES, internal ribosome entry site; pA, polyadenylation signal; Blast, Blasticidin resistance. (B) Brightfield and green fluorescence images of virus-producing HEK293 cells transfected with either linGFP-pLenti or spGFP-pLenti. Faint GFP positivity is evident in cells transfected with spGFP-pLenti, indicating backsplicing. (C) qRT-PCR showing the expression of circular spGFP (left panel) and linear/circular GFP (right panel) in virus-producing HEK293 cells transfected with either linGFP-pLenti or spGFP-pLenti. Each dot represents one biological replicate. (D) Western blot showing the expression of GFP in virus-producing HEK293 cells transfected with either linGFP-pLenti or spGFP-pLenti. (E) Viral titer produced by HEK293 cells transfected with either linGFP-pLenti or spGFP-pLenti measured with Takara’s Lenti-X GoStix kit. (F) Infection of 501Mel and WM164 human melanoma cell lines with linGFP-pLenti or spGFP-pLenti viral supernatants. Infected cells were selected with 10 µg/mL Blasticidin for 1 week and then stained with Crystal Violet (left panels). The quantification of the extracted dye is shown in the right panels. (G) Brightfield and green fluorescence images of Blasticidin-selected 501Mel and WM164 cells infected with linGFP-pLenti or spGFP-pLenti viral supernatants. (H) qRT-PCR showing the expression of linear/circular GFP and circular spGFP in 501Mel and WM164 cells infected with linGFP-pLenti or spGFP-pLenti viral supernatants. Expression of spGFP was not detected (ND) with divergent primers. Uninfected parental cell lines are included as GFP-negative controls. ****, p<0.0001

### Stable circRNA overexpression using a transposon

Backsplicing of the viral transcript in virus-producing HEK293 cells may interfere with the production of viral particles carrying an intact, non-spliced circRNA transgene. We therefore decided to test a non-viral circRNA transgene delivery method using a transposon. We modified the ATP1 transposon^31^ harboring inverted repeats for the Sleeping Beauty and piggyBac transposases to contain the spGFP reporter under the control of the EF1α promoter and a Blasticidin selection cassette (Fig. 2A). We transfected HEK293 cells with the spGFP transposon along with expression plasmids encoding the Sleeping Beauty or piggyBac transposases. Both combinations resulted in robust expression of circularized spGFP, as determined by qRT-PCR with divergent primers (Fig. 2B). In addition, we observed expression of GFP by Western blot in these cells (Fig. 2C). Next, we tested whether circular spGFP expression is maintained over multiple passages, which would indicate that copies of the spGFP transposon are integrated into the genome. To this end, we transfected HEK293 cells with the spGFP transposon and either Sleeping Beauty or piggyBac expression vectors or empty pCDNA3 and determined the expression of circular spGFP at 2 and 10 passages post-transfection. Both in the presence and absence of Blasticidin, circular spGFP expression declined after 10 passages (Fig. 2D). However, in the presence of Sleeping Beauty and especially piggyBac, spGFP levels were maintained at higher levels compared to the empty pCDNA3-transfected control (Fig. 2D). We next tested whether a similar effect is observed in human melanoma cells by transfecting A375 cells with the spGFP transposon and either Sleeping Beauty or piggyBac expression vectors or empty pCDNA3. The transfected A375 cells were cultured for 10 passages in the absence of Blasticidin and then placed on Blasticidin selection. Significantly more cells survived selection when Sleeping Beauty or piggyBac were expressed (Fig. 2E), suggesting the transposon was maintained long-term in the absence of selection. To test whether transposon maintenance was due to genomic integration, we cloned Blasticidin-selected single cell A375 clones that were transfected with the spGFP transposon and piggyBac expression plasmid and isolated genomic DNA. We digested the DNA with MspI to cut within the transposon and in juxtaposed genomic regions and subjected the DNA to Southern blot analysis using a Blasticidin probe. This strategy detected multiple bands of different sizes in most clones (Fig. 2F), validating that the transposon was integrated in the genome. Finally, to confirm that the transposon is amenable for delivering other circRNA transgenes, we replaced the spGFP construct with the exons of the human circDNAJC2 and circPMS1 circular RNAs (Fig. 2G). We delivered these circRNA transposon constructs along with piggyBac to WM164 cells and selected in Blasticidin. Notably, this resulted in significant overexpression of circDNAJC2 and circPMS1 as determined by qRT-PCR using divergent primers (Fig. 2G). Thus, stable circRNA overexpression is readily achieved with a transposon as delivery vehicle.

**Figure 2:**
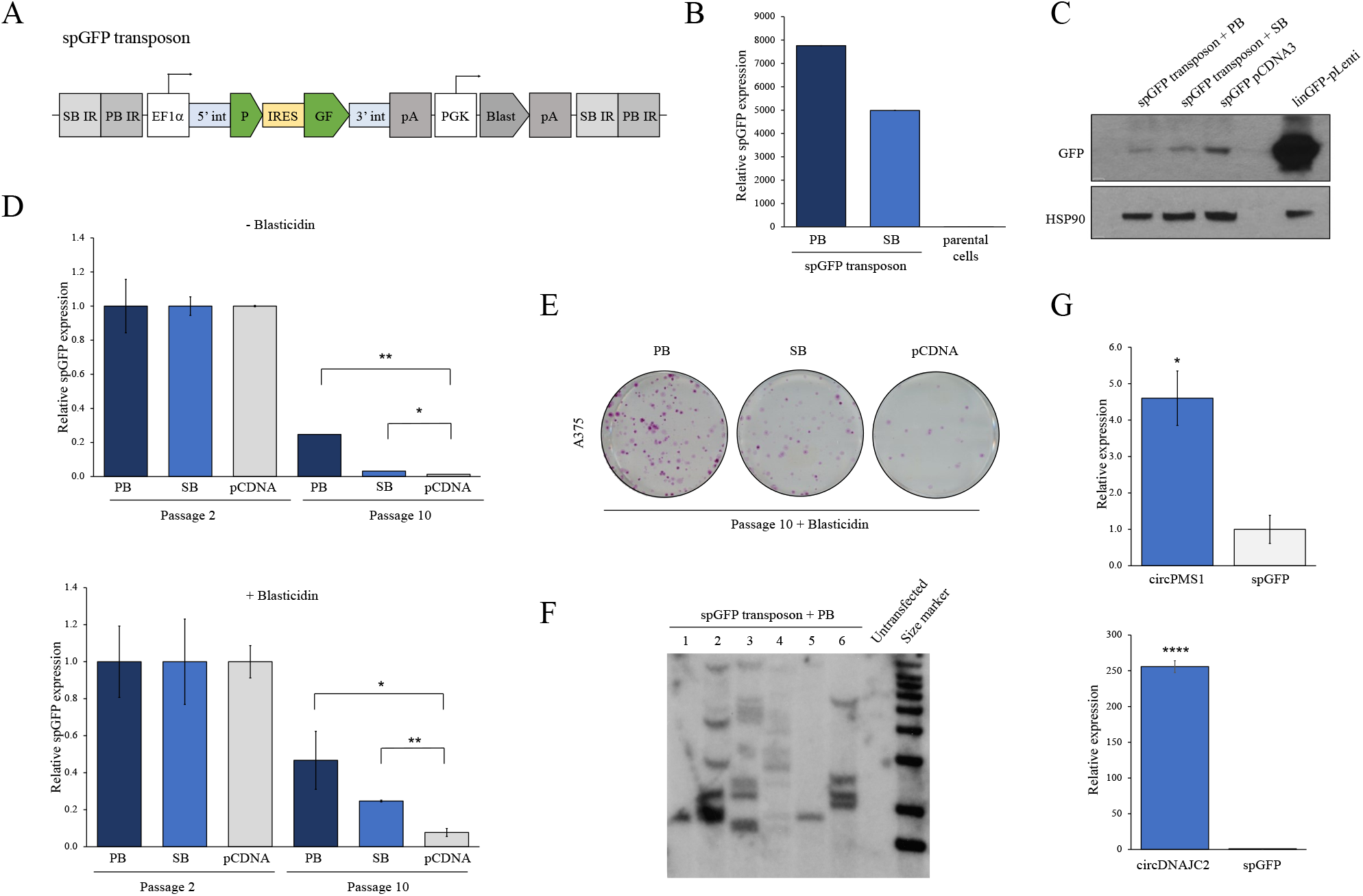
circRNA construct delivery and stable expression with a transposon. (A) Schematic outline of the spGFP transposon. SB IR, inverted repeat for Sleeping Beauty transposase; PB IR, inverted repeat for piggyBac transposase; 5’int, human ZKSCAN1 5’ intron; 3’int, human ZKSCAN1 3’ intron; IRES, internal ribosome entry site; pA, polyadenylation signal; Blast, Blasticidin resistance. (B) qRT-PCR showing expression of circular spGFP in HEK293 cells transfected with the spGFP transposon and either piggyBac (PB) or Sleeping Beauty (SB) expression vectors. Untransfected parental cells are included as negative controls. (C) Western blot showing expression of GFP in HEK293 cells transfected with the spGFP transposon and either piggyBac (PB) or Sleeping Beauty (SB) expression vectors. A circular spGFP expression plasmid (spGFP-pCDNA3) and a linear GFP (linGFP-pLenti) lentiviral vector were transfected into HEK293 and used as controls for GFP expression. (D) qRT-PCR showing circular spGFP expression in HEK293 cells transfected with the spGFP transposon and either piggyBac (PB) or Sleeping Beauty (SB) or empty pCDNA3. Circular spGFP expression was analyzed 2 and 10 passages after transfection in either unselected cells (top panel) or cells kept on 10µg/mL Blasticidin (bottom panel). (E) Crystal Violet staining of human A375 melanoma cells transfected with the spGFP transposon and either piggyBac (PB) or Sleeping Beauty (SB) or empty pCDNA3. Cells were cultured for 10 passages and then selected in 10 µg/mL Blasticidin. (F) Southern blot demonstrating the integration of the spGFP transposon in A375 clones transfected with spGFP transposon vector and a piggyBac expression plasmid. (G) Expression of circPMS1 and circDNAJC2 in WM164 melanoma cells transfected with either circPMS1 transposon (top panel) or circDNAJC2 (bottom panel) transposon and a piggyBac expression plasmid. *, p<0.05; **, p<0.01; ****, p<0.0001.

### In vivo delivery of a circRNA construct using transposons

Following the establishment of a circRNA transposon expression system in cultured cells, we assessed whether the transposon approach is suitable for delivering circRNA overexpression constructs *in vivo*. Specifically, we examined whether a circRNA construct can be delivered to the liver of mice via hydrodynamic tail vein injection. To enrich for cells that contain the circRNA construct, we used a hepatocellular carcinoma model in which tumorigenesis is driven by delivering a Sleeping Beauty transposon harboring a c-Myc cDNA^32^. We combined the spGFP transposon plasmid with a Sleeping Beauty transposase expression plasmid, the c-Myc transposon, and a plasmid encoding Cas9 and a sgRNA targeting p53^32^ and performed hydrodynamic tail vein injections in wildtype C57BL/6 mice (Fig. 3A). After aging the mice for 4 weeks we isolated the livers and out of the 5 mice injected, 2 mice had developed hepatocellular carcinomas. We first isolated genomic DNA from 4 liver tumors and performed qPCRs using a primer pair designed over the Blasticidin cassette. This demonstrated that the spGFP transposon DNA was present in the tumors (Fig. 3B). Next, we isolated RNA from the same liver tumors and performed a qRT-PCR for circular spGFP using divergent primers. All tumors tested exhibited expression of circular spGFP, albeit at varying levels (Fig. 3C). These data demonstrate that circRNAs can be ectopically expressed in mouse liver by delivering circRNA transgene-containing transposons via hydrodynamic tail vein injection.

**Figure 3:**
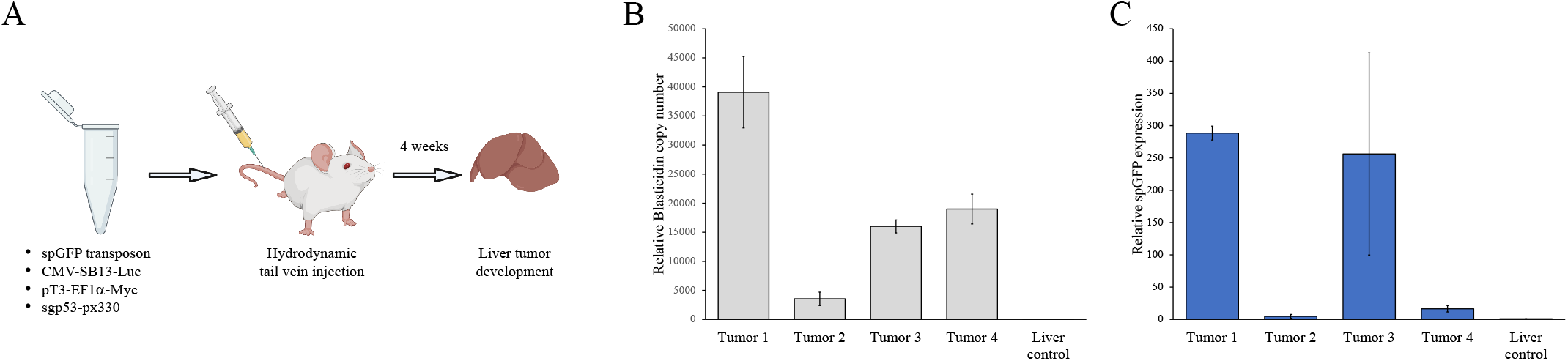
Ectopic circRNA expression in a hepatocellular carcinoma model with transposon-based delivery. (A) Outline of the hydrodynamic tail vein injection to generate a hepatocellular carcinoma model. (B) qPCR copy number analysis on genomic DNA isolated from liver tumors to detect Blasticidin as a surrogate for the presence of the spGFP transposon. Normal liver from uninjected mice was used as negative control. (C) qRT-PCR analysis on RNA isolated from liver tumors using divergent primers to quantify the expression of circular spGFP. Normal liver from uninjected mice was used as negative control.

### Global circRNA overexpression in genetically engineered mice

While the *in vivo* delivery of transposon DNA is readily achievable in livers, it is far more difficult to accomplish in other organs and cell types. To study gene functions *in vivo*, overexpression constructs are typically inserted into the mouse germline of genetically engineered mice. However, to our knowledge this approach has not been used for *in vivo* overexpression of circRNAs. Thus, we tested whether an ectopic circRNA knock-in allele is functional in mice. To this end, we cloned a targeting vector for recombination-mediated cassette exchange (RMCE) into embryonic stem cells (ESCs) harboring a homing cassette (CHC) downstream of the collagen1a1 locus (Fig. 4A). This targeting vector contains the spGFP circRNA reporter under the control of the constitutive EF1α promoter as well as a FRT site and a PGK promoter to render it compatible with RMCE (Fig 4B). We targeted C10 v6.5 ESCs^33^ with this spGFP expression construct and validated correct transgene integration by PCR. We then injected one positively targeted ESC clone into Balb/c blastocysts followed by transfer into pseudopregnant CD1 females. Three chimeras were born with ESC contribution based on coat-color ranging from 25-80% (Fig. 4C). We euthanized the chimeras at 4 weeks of age and collected all major organs, isolated RNA, and performed qRT-PCR to detect spGFP expression with divergent primers. Notably, spGFP expression was detected in all organs (Fig. 4D). Thus, using a ubiquitous promoter and the human ZKSCAN1 introns enables expression of circRNAs in various organs of knock-in mice.

**Figure 4:**
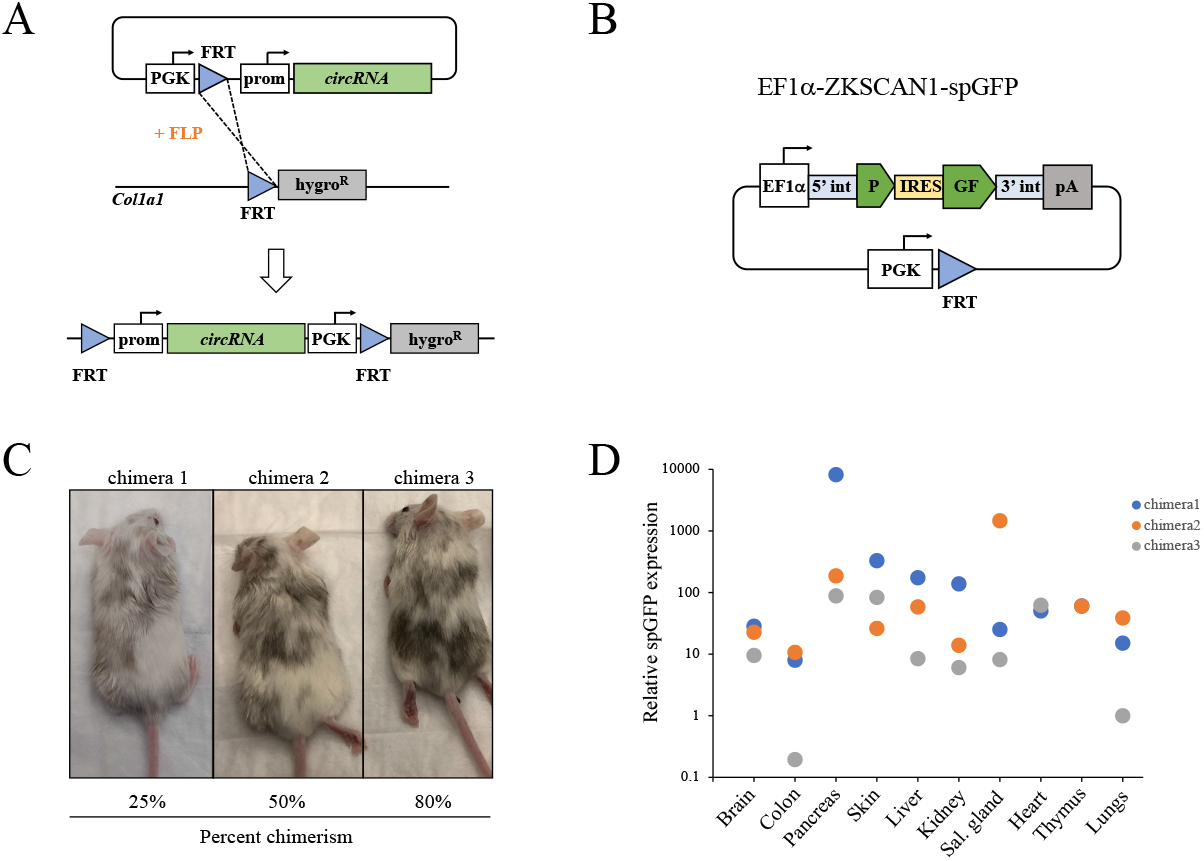
Global ectopic circRNA expression in genetically engineered knock-in mice. (A) Schematic outline of the recombination-mediated cassette exchange (RMCE) approach. (B) Schematic outline of the constitutive and ubiquitous circular spGFP targeting vector (EF1α-ZKSCAN1-spGFP). 5’int, human ZKSCAN1 5’ intron; 3’int, human ZKSCAN1 3’ intron; IRES, internal ribosome entry site; pA, polyadenylation signal. (C) Images of the three EF1α-ZKSCAN1-spGFP chimeras. The estimated percentage of chimerism based on coat color is indicated. (D) qRT-PCR showing circular spGFP expression in organs from EF1α-ZKSCAN1-spGFP chimeras. Sal. gland, salivary gland.

### Tissue-specific and inducible circRNA expression in a mouse melanoma model

Having achieved global expression of the spGFP reporter circRNA in mice, we assessed whether ectopic circRNA expression can be induced and directed to a specific cell type *in vivo*. To this end, we used our ESC-genetically engineered mouse modeling (ESC-GEMM) melanoma platform^34^ consisting of ESCs derived from GEMMs harboring alleles to efficiently induce melanoma formation, integrate transgenes by RMCE, and control transgene expression with the Tet-ON system. Specifically, we used the BPP model, which harbors LSL-Braf^V600E^ and Pten^FL/FL^ driver alleles, a melanocyte-specific, 4OH-Tamoxifen (4OHT)-inducible Tyr-CreERt2 recombinase allele, the Cre-dependent CAGs-LSL-rtTA3 transactivator, and the CHC homing allele. When chimeric mice generated with the BPP model are topically treated with 4OHT on their back skin, melanomas form within 6 weeks. Moreover, expression of inducible transgenes is activated specifically in melanoma cells by switching chimeras to a Dox-containing diet. This model is thus ideal to rapidly test if ectopic circRNA expression can be directed to melanoma cells.

To generate an inducible spGFP circRNA reporter targeting vector, we replaced the EF1α promoter with a Doxycycline-inducible TRE promoter (Fig. 5A). We targeted BPP ESCs with the TRE-spGFP construct by RMCE and produced chimeric mice via blastocyst injection (Fig. 5B). At weaning, we shaved the chimeras’ back skin with clippers and topically applied either several 1µL drops of 25mg/mL 4OHT in DMSO or enough 4OHT solution (2.5mg/mL) to cover the back skin. After 5-6 weeks when melanomas had formed, we switched chimeras from a regular diet to a 200mg/kg Dox diet for one week. We then harvested melanomas (Fig. 5C) and isolated RNA. As there is no selective pressure to recombine the CAGs-LSL-rtTA3 allele in contexts where the inserted transgene does not accelerate melanomagenesis^34^, we first validated CAGs-LSL-rtTA3 recombination. To do so, we assessed expression of rtTA3 in melanomas by qRT-PCR. Of the 9 isolated tumors, 5 had the CAGs-LSL-rtTA3 allele recombined and expressed rtTA3, while 4 tumors did not (Fig. 5D). We then determined if circular spGFP was expressed in melanomas by qRT-PCR using divergent primers. Notably, circular spGFP was detected in Dox-treated tumors that also express rtTA3 (Fig. 5E) while circular spGFP expression was not detected in Dox-treated tumors that lacked rtTA3 expression or in livers (Fig. 5E). Thus, ectopic circRNA expression can be spatiotemporally controlled *in vivo* by using the appropriate Cre strain and the Tet-ON system.

**Figure 5:**
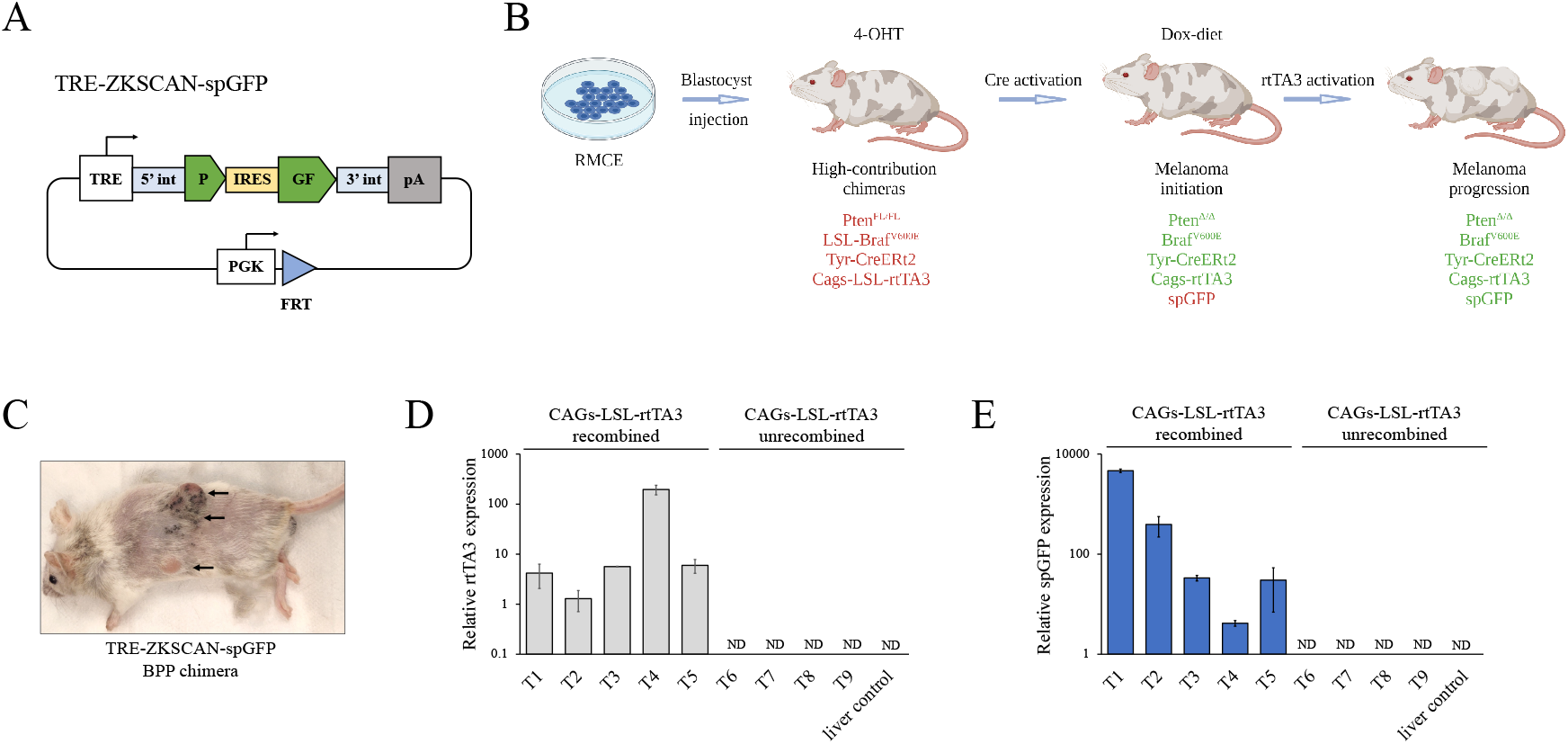
Melanoma cell-specific ectopic circRNA expression in a genetically engineered melanoma mouse model. (A) Schematic outline of the Dox-inducible circular spGFP targeting vector (TRE-ZKSCAN1-spGFP). TRE, Tetracycline-response element promoter; 5’int, human ZKSCAN1 5’ intron; 3’int, human ZKSCAN1 3’ intron; IRES, internal ribosome entry site; pA, polyadenylation signal. (B) Outline of the ESC-GEMM approach. BPP ES cells are targeted with the circRNA expression construct by RMCE. Targeted ES cells are used to generate chimeras, which are treated with 4-OH Tamoxifen (4-OHT) to activate melanocyte-specific Cre recombinase. Cre induces expression of Braf^V600E^ and rtTA3 and deletes Pten. Subsequent administration of Doxycycline activates the Tet-ON system and induces expression of the circRNA expression construct. (C) Image of a BPP chimera harboring the TRE-ZKSCAN1-spGFP construct. Melanomas are indicated by arrows. (D) qRT-PCR showing the expression of rtTA3 in melanomas. Tumors in which the CAGs-LSL-rtTA3 allele is recombined express the rtTA3 transactivator. (E) qRT-PCR showing the expression of circular spGFP in melanomas. Melanomas that express rtTA3 also express spGFP. ND, not detected.

## Discussion

In this work we established new genetic tools for the stable, long-term overexpression of candidate circRNAs in cultured mammalian cells *in vitro* and in mice. We used a transposon approach to deliver a circRNA expression cassette to cell lines, where it is integrated into the genome upon transient co-expression of a transposase. We further show that circRNA expression cassettes can be delivered to the livers of mice via hydrodynamic tail vein injections. In addition, using a recombination-mediated cassette exchange approach, we inserted a circRNA expression cassette in the genome of mouse ESCs, enabling the generation of mice that ubiquitously express an ectopic circRNA. We used the same approach to generate a melanoma mouse model in which ectopic circRNA expression can be induced specifically in melanoma cells. These tools and approaches will pave the way for functional analyses of circRNA function and overcome some of the limitations of current circRNA expression systems. In particular, we are unaware of any attempts to overexpress circRNAs *in vivo* and our work demonstrates the feasibility of generating circRNA transgenic mice. This will stimulate further research of circRNA function in normal physiology and disease.

We observed that circRNA expression cassettes in lentiviral constructs are backspliced in the virus-producing cells. This may interfere with packaging the viral transcript into viral particles. Moreover, viral transcripts that have undergone backsplicing and are packaged into viral particles would lack the circRNA expression cassette. These considerations offer possible explanations as to why we observed a high viral titer with the spGFP-pLenti lentiviral construct but failed to obtain appreciable Blasticidin resistance or circular spGFP expression in melanoma cells infected with the viral supernatant. The extent of backsplicing in virus-producing HEK293 cells is likely dictated, at least in part, by the introns used to mediate circularization. The human ZKSCAN1 introns used in our construct efficiently promote circularization^29^, which in this case is a detriment. The ZKSCAN1 introns could be replaced by introns that are less efficient at mediating circularization to prevent backsplicing in HEK293 cells. However, this would also reduce the extent of circularization in infected cells and increase the abundance of unspliced linear viral transcript containing the circRNA exons and flanking introns. It may therefore be challenging to discern whether any observed phenotype is elicited by the circRNA or the linear transcript. One could use RNAi or inhibitors targeting the splicing machinery to transiently reduce splicing in virus-producing cells. However, this may negatively affect HEK293 cells and thereby lower the quality of the produced virus. We found the transposon approach to be an easy-to-use alternative to lentiviruses for stable circRNA overexpression.

Given that the transposon is not delivered to target cells via an RNA intermediate, this approach is not hindered by the shortcomings of lentiviral circRNA delivery. However, as the transposon and transposase expression plasmids are co-transfected, this approach may be limited in difficult-to-transfect cell types. Nevertheless, we observed long-term expression of the circular spGFP reporter in melanoma cell lines using two different transposases, Sleeping Beauty and piggyBac. The modular nature of the transposon allows for the easy exchange of the promoter, the circularizing introns, or the selection marker, allowing the generation of custom-built transposons that fit one’s need. One may also wish to optimize the DNA delivery and transposon:transposase plasmid ratios to normalize transposon integration in a polyclonal cell population. As was evident from our transposon genomic DNA integration analysis, individual subclones had dissimilar numbers of transposons integrated. This would result in varying expression levels of the candidate circRNA in different subclones, which may influence the observed phenotype if subclones enrich or drop out over time. Selecting a few individual subclones with the desired extent of circRNA overexpression for downstream phenotypic analyses is therefore recommended.

We successfully generated genetically engineered mice in which an ectopic circRNA was expressed constitutively in all cells or inducibly in melanocytes/melanoma cells. Given that ectopic circRNA expression was possible with both approaches, we surmise that most, if not all, gene targeting approaches to produce overexpression mouse strains are compatible with circRNA expression constructs. circRNA depletion via genetic knock-out, as has been done for CDR1as^35^, is feasible and useful for transcripts that are primarily circularized like CDR1as^3,36^. However, most circRNAs are encoded by genes that also encode protein-coding mRNAs. The knock-out of circRNA exons in those genes will therefore also affect the encoded protein, which complicates the analysis of any observed phenotypes. Sophisticated methods that target the flanking introns to prevent circularization without affecting linear splicing, for instance by CRISPR-Cas9-mediated Alu site deletion, await testing and their success is likely going to be locus dependent. Thus, circRNA overexpression in genetically engineered mice to study circRNA function *in vivo* is currently the more practical approach. In summary, our tools and approaches expand our genetic toolkit to overexpress circRNAs *in vitro* and *in vivo* and will stimulate studies to functionally characterize circRNAs.

## Methods

### Plasmids and cloning

To generate spGFP-pLenti, pcDNA3-ZKSCAN1-spGFP (Addgene) was digested with BamHI and XhoI and the spGFP-ZKSCAN1 cassette ligated into BamHI/SalI-digested pLenti-EF1α-GFP_miRE to replace GFP_miRE. The EF1α-spGFP-ZKSCAN1-Blast transposon was generated by digesting the ATP1 transposon (provided by Roland Rad) with BstXI to remove the CAGS promoter, polyadenylation signals, and splice acceptor/donor sites and replace them with a multiple cloning site containing MluI-NotI-BglII sites by oligo cloning. A SV40 polyA-PGK promoter-BlasticidinR cDNA-bGH polyA cassette was then synthesized by GeneArt (Thermo Fisher), PCR amplified, and InFusion (Clonetech/Takara) cloned into NotI/BglII digested ATP1-MCS, generating ATP1-Blast. Finally, EF1α-spGFP-ZKSCAN1 was PCR amplified from spGFP-pLenti and InFusion cloned into MluI/NotI digested ATP1-Blast. To generate the EF1α-spGFP targeting vector, a EF1α-LSL-GFP targeting vector was digested with XhoI/EcoRI. The spGFP-ZKSCAN1 cassette was PCR amplified from spGFP-pLenti to introduce restriction sites, the PCR product was digested with XhoI/EcoRI, and ligated into the targeting vector. To generate the TRE-spGFP targeting vector, the spGFP-ZKSCAN1 cassette was PCR amplified and InFusion cloned into XhoI/EcoRI digested cTRE-CHC targeting vector (provided by Lukas Dow). The circPMS1 and circDNAJC2 transposons were cloned by synthesizing the circRNA exons by GeneArt, followed by PCR amplification and InFusion cloning into SnaBI/PstI digested EF1α-spGFP-ZKSCAN1-Blast transposon plasmid.

### Cell culture

Human melanoma cell lines were cultured in RPMI-1640 (Lonza) containing 5% FBS and HEK293T-LentiX in high-glucose DMEM (Sigma) containing 10% FBS in a 5% CO2 incubator at 37°C. For Crystal Violet staining, cells were washed in phosphate-buffered saline (PBS) and stained with 0.1% Crystal Violet (VWR) in 20% methanol. The dye was extracted with 100 µL of acetic acid and absorbance was measured at 600 nm.

### Transfection and virus production

For transposon transfection, 0.9 µg of spGFP transposon and 0.1 µg of either pCMV-HSB5 (Sleeping Beauty transposase), pCDNA-mPB (piggyBac transposase), or empty pCDNA3.1 were transfected into HEK293T-LentiX or melanoma cells at 80-90% confluence using JetPrime (Polyplus) according to the manufacturer’s recommendations. pCMV-HSB5 and pCDNA-mPB were provided by David Adams. Where indicated, transfected cells were selected with 10 µg/mL Blasticidin for 10 days starting 24 hours following the transfection. For lentivirus production, HEK293T-LentiX cells were transfected at 90% confluence with 6 µg of lentiviral vector, 0.66 µg of VSVG envelope vector, and 5.33 µg of Δ8.2 packaging vector using JetPrime according to the manufacturer’s recommendations. Viral supernatant was collected after 48 hours, filtered through 0.45µm syringe filters, and used to infect melanoma cells in the presence of 8 µg/mL polybrene. Viral titer was determined using the Lenti-X GoStix kit (Takara) following the manufacturer’s recommendations

### RNA isolation, cDNA synthesis, and quantitative real-time PCR

For RNA isolation from cell lines, cells were washed with PBS and collected by scraping and centrifugation. Cell pellets were resuspended in TRI reagent (Zymo Research) and RNA was isolated as per the manufacturer’s instructions. For RNA isolation from organs and tumors, tissue pieces were physically disrupted in 600 µL of TRI reagent in high-impact zirconium beads (Benchmark) using a microtube homogenizer (BeadBug). RNA was then isolated from lysate diluted 1:2-1:3 in TRI reagent according to the manufacturer’s recommendations. RNA was treated with RQ1 RNase-free DNase (Promega) following the manufacturer’s instructions. cDNA was synthesized using the 5X PrimeScript RT Master Mix (Clontech/Takara), following the manufacturer’s instructions. qRT-PCRs were performed using PerfeCTa SYBR Green FastMix (Quantabio) and results were analyzed using the ΔΔCt method and Actin was used as a normalization control. The following primers were used: spGFP forward 5’-ATGGCAACATCCTGGGCAAT -3’, reverse 5’-TTCACATCGCCATTCAGCTC -3’; circPMS1 forward 5’-TCCAAGATCTCCTCATGAGC-3’, reverse 5’-TACAACACTGACCACCGAAG-3’; circDNAJC2 forward 5’-AGAAGCTGCTCGGTTAGCTA-3’, reverse 5’-GTGCTCTTGTTGCTCTGTTC-3’; Actin forward 5’-TTGCTGACAGGATGCAGAAG -3’, reverse 5’-ACATCTGCTGGAAGGTGGAC-3’.

### Western blot

Cells were lysed in RIPA buffer and 20 μg of protein lysate were separated on NuPAGE 4-12% Bis-Tris Midi Gels (Invitrogen). Proteins were transferred onto nitrocellulose membrane and stained with Ponceau Red to confirm complete transfer. Membranes were blocked in 5% non-fat dry milk in TBS-T and incubated over night at 4°C with anti-GFP (1:1,000; Cell Signaling Technologies) or anti-Actin (1:4,000; Invitrogen) antibodies diluted in 5% non-fat dry milk in TBS-T. The membranes were washed with TBS-T and incubated with HRP-conjugated secondary anti-rabbit antibody (1:20,000; Jackson ImmunoResearch) for 1 hour. The signal was detected with SuperSignal West Pico PLUS Chemoluminescent Substrate (Thermo Scientific) following the manufacturer’s instructions.

### Southern blot

A375 cells were transfected with 0.9 µg of spGFP transposon and 0.1 µg of pCDNA-mPB (piggyBac transposase) and selected with 10 µg/mL Blasticidin for 3 weeks. Clones derived from single cells were picked using cloning cylinders and expanded. Genomic DNA was isolated with Proteinase K lysis buffer and 15 µg of DNA were digested overnight with 40 units of MspI (New England Biolabs). Digested DNA was separated on a 0.8% agarose gel, followed by in-gel depurination, denaturation, and neutralization. DNA was capillary transferred in 10xSSC onto nitrocellulose membrane and crosslinked by baking at 70°C for one hour. A Blasticidin probe was PCR amplified, labeled with α-^32^P-dCTP using DECAprime II random prime labeling kit (Invitrogen) and purified with MicroSpin G-50 columns (Cytiva) using the manufacturers’ recommendations. The membrane was incubated with labeled probe and 250 µg/mL salmon sperm DNA (Roche) at 65°C overnight in Perfecthyb Plus Hybridization Buffer (Millipore), washed in 2xSSC plus 0.5% SDS, and exposed to X-ray film for 4 days.

### Embryonic stem cell targeting and mouse generation

C10 v6.5 ES cells were obtained from Rudolf Jaenisch. BPP ES cells were generated in our laboratory^34^. ES cells were electroporated with 15 µg targeting vector and 7.5 µg pCAGGS-FLPe and selected with 125 µg/mL hygromycin. Clones were picked after 7-10 days, expanded, and verified by PCR genotyping of the targeted allele. Targeted ES cells were injected into Balb/c blastocysts and transferred into pseudopregnant CD1 females. ES cell targeting and blastocyst injections were performed by the Moffitt Gene Targeting Core.

### Mouse husbandry and hydrodynamic tail vein injection

All animal experiments were conducted in accordance with an IACUC protocol (R-IS00005420) approved by the University of South Florida. For hydrodynamic tail vein injection, C57Bl/6 mice (Jackson Laboratory) were injected with a volume of sterile 0.9% saline solution equivalent to 10% of their body weight. The saline solution contained 10 µg spGFP transposon plasmid, 2.5 µg CMV-SB13-Luc, 10 µg pT3-EF1a-Myc, 10 µg sgp53-px330.

### Statistical analysis

Statistical analysis was performed using the unpaired two-tailed t-test. All in vitro experiments were performed with three biological and three technical replicates. Unless otherwise noted, the mean ± SEM of one representative experiment is shown. A p-value below 0.05 was considered statistically significant.

## Acknowledgements

We thank the members of the Karreth lab for critical reading of the manuscript, Dr. Roland Rad and David Adams for plasmids, and Dr. Rudolf Jaenisch for ES cells. Figures 3A and 5B were created with BioRender.com.

## Author Contribution

Conceptualization, FAK; Methodology, NM, AN, OV, and FAK; Investigation, NM, AN, OV, AF, and FAK; Writing – Original Draft, FAK; Writing – Reviewing and Editing, NM, OV, AN, AF, GMD, and FAK; Supervision, GMD and FAK.

